# Within-host rhinovirus evolution in upper and lower respiratory tract highlights capsid variability and mutation-independent compartmentalization

**DOI:** 10.1101/2023.05.11.540440

**Authors:** Negar Makhsous, Stephanie Goya, Carlos Avendaño, Jason Rupp, Jane Kuypers, Keith R. Jerome, Michael Boeckh, Alpana Waghmare, Alexander L Greninger

**Author notes:** The authors contributed equally to this manuscript (Negar Makhsous, Stephanie Goya, Carlos Avendaño). ^#^Corresponding authors, **Corresponding author contact** Alexander L Greninger, 1616 Eastlake Ave East Suite 320, Seattle, WA 98102, Alpana Waghmare, 1100 Eastlake Ave East, E4-100, Seattle, WA 98109.

## Abstract

**Background:** Human rhinovirus (HRV) infections can progress from the upper (URT) to lower (LRT) respiratory tract in immunocompromised individuals, causing high rates of fatal pneumonia. Little is known about how HRV evolves within hosts during infection.

**Methods:** We sequenced HRV complete genomes from 12 hematopoietic cell transplant patients with prolonged infection for up to 190 days from both URT (nasal wash, NW) and LRT (bronchoalveolar lavage, BAL) specimens. Metagenomic (mNGS) and amplicon-based NGS were used to study the emergence and evolution of intra-host single nucleotide variants (iSNVs).

**Results:** Identical HRV intra-host populations in matched NW and BAL specimens indicated no genetic adaptation is required for HRV to progress from URT to LRT. Microbial composition between matched NW and BAL confirmed no cross-contamination during sampling procedure. Coding iSNVs were 2.3-fold more prevalent in capsid over non-structural genes, adjusted for length. iSNVs modeled onto HRV capsid structures were significantly more likely to be found in surface residues, but were not preferentially located in known HRV neutralizing antibody epitopes. Newly emergent, serotype-matched iSNV haplotypes from immunocompromised individuals from 2008-2010 could be detected in Seattle-area community HRV sequences from 2020-2021.

**Conclusion:** HRV infections in immunocompromised hosts can progress from URT to LRT with no specific evolutionary requirement. Capsid proteins carry the highest variability and emergent mutations can be detected in other, including future, HRV sequences.

## Introduction

Human rhinoviruses (HRVs) constitute the most prevalent cause of the common cold but are also associated with exacerbations of pre-existing airway diseases such as asthma and COPD [1–3]. HRV infection progression from the upper (URT) to lower respiratory tract (LRT) is well-described in hematopoietic cell transplant (HCT) recipients [4–6]. Once LRT infection has developed, rates of HRV-associated respiratory failure and fatal pneumonia are higher among HCT recipients and are comparable to those seen with other respiratory viruses including respiratory syncytial virus, parainfluenza virus, and influenza virus [7,8].

HRVs belong to the *Picornaviridae* family and are divided into HRV-A, HRV-B, and HRV-C species, which are themselves divided into serotypes based on either neutralization assays or genotyping. HRVs consist of a positive-sense single-stranded RNA genome of approximately 7000 nt length that is organized in a single ORF that generates four capsid proteins (VP1-4) and seven non-structural proteins (2A-2C and 3A-3D). The VP1 nucleotide region serves as the basis for phylogenetic serotype classification of HRVs [1,9]. A total of 164 HRV serotypes have been identified (79 HRV-A, 30 HRV-B, and 55 HRV-C types) which can co-circulate throughout the year, complicating both vaccine and drug development [2,10–13].

While studies have focused on HRV species and serotype surveillance, there is scarce knowledge about how HRV diversity is triggered within hosts and how the virus progresses from the URT to LRT in immunocompromised individuals [14–17]. In this study, we analyzed within-host HRV evolution dynamics in 12 HCT recipients with prolonged infection by different HRV species, comparing both URT and LRT specimens.

## Materials and Methods

### Patient and viral load testing

URT (nasal wash, NW) and LRT (bronchioalveolar lavage, BAL) samples were collected from allogeneic HCT recipients after being screened by RT-qPCR during the routine follow up at the UW Medicine/Fred Hutchinson Cancer Center from 2006 to 2014. This study was approved by the Institutional Review Board at the Fred Hutchinson Cancer Center. Subjects signed informed consent permitting the use of data and samples for research.

### Genome sequencing and phylogenetic analysis

Genome sequencing and analysis methods are more extensively discussed in Supplemental Material. Briefly, metagenomic next-generation sequencing (mNGS) libraries were prepared as described previously [18,19]. In addition, 1kb tiling amplicon-based sequencing libraries were used to obtain coverage in specimens where mNGS failed. Consensus genomes were called using the Revica pipeline (https://github.com/greninger-lab/revica) using a database containing all existing rhinovirus complete genomes from ICTV [11]. Maximum likelihood trees were inferred with IQ-TREE v2.1 using SH-aLRT test (1000 replicates) and UFBoot2 method (10000 replicates) to evaluate reliability of sequence clusters [20–22].

### Intra-host single nucleotide variant (iSNV) analysis

iSNVs were called using the LAVA pipeline [23] using bwa-mem [24] with a minimum allele frequency (MAF) of 10% in the positions with a minimum sequence depth of coverage of 30X. All iSNVs are listed in Supplemental Table 1. Microbiome composition for URT and LRT specimens with mNGS data was assessed using Kraken2 software compared to the standard prebuilt genomes database (v20221209) [25] and the phyloseq library in RStudio [26] and plotted with ggplot2 library [27]. Capsid iSNVs were mapped onto the 3D viral capsid structure of HRV-A16 (PDB: 1AYM) using HHPred and visualized using Pymol (Supplemental Table 2) [28,29]. HHPred was also used to map iSNVs onto the Coxsackievirus B3 capsid (PDB: 4GB3) and correlated with published mutational fitness effect (MFE) measures using Mann Whitney Wilcoxon statistical test [30].

## RESULTS

### HRV genome sequencing and viral serotype assignment

A total of 12 HCT recipients with HRV positive specimens from 2006 to 2014 were included (Table 1). Except for a 1-year-old, all subjects were adults (age range: 24 – 67 years-old, median: 46 years-old). Ten of the twelve individuals had low absolute lymphocyte count (<1000 cells/µL) and all had respiratory symptoms at the time of the initial HRV infection detection (Table 1). In addition, all individuals developed proven or probable lower respiratory tract infection, considered as HRV detected in BAL despite the presence or absence of radiographic abnormalities. Ten of the twelve individuals died in the 5 months following the last HRV positive detection (except individuals S01 and S06).

**Table 1.** Epidemiological information summary. The age and sex of each individual is detailed as well as the HRV serotype detected. The time from the hematopoietic cell transplantation (HCT) to the first HRV positive sample detected is informed in days. The aboslute lymphocyte count was evaluated at the time of the first HRV positive sample detected. HRV human rhinovirus; HCT, hematopoietic cell transplantation.

| Individual | Age<br>(years) | Sex | HRV<br>Serotype | Time from HCT to HRV<br>infection (days) | Absolute lymphocyte<br>count (k/μL) |
| --- | --- | --- | --- | --- | --- |
| S01 | 46 | Male | C17 | 88 | 0.68 |
| S02 | 1 | Male | C36 | 273 | 0.55 |
| S03 | 50 | Male | B97 | 0 | 0.05 |
| S04 | 50 | Female | B06 | 246 | 0.57 |
| S05 | 64 | Male | A57 | 36 | 0.18 |
| S06 | 34 | Female | A105 | 5 | 1.49 |
| S07 | 67 | Female | A58 | 414 | 2.34 |
| S08 | 37 | Male | A78 | 162 | 0.50 |
| S09 | 24 | Male | A102 | 18 | 0.11 |
| S10 | 41 | Male | A39 | 181 | 0.11 |
| S11 | 63 | Male | A82 | 14 | 0.59 |
| S12 | 44 | Male | C28 | 108 | 0.00 |

RT-qPCR Ct values were variable among samples and individuals (range: 17.5 – 34.3). mNGS failed to recover genomes in seven specimens, which required amplicon-based sequencing for genome recovery (Table 2). Comparison of mNGS and amplicon-based sequencing methods in select samples showed high levels of consistency in variant allele frequencies recovered (r^2^>0.96, Supplemental Figure 1). mNGS analysis revealed a coinfection with HCoV 229E and metapneumovirus in BAL10b.

**Table 2.** HRV sequencing and intra-single nucleotide variant (iSNV) characterization. For each individual, the upper (nasal wash, NW) and lower (bronchioalveolar lavage, BAL) sample is identified and the collection date, the days elapsed since the first HRV positive and the RT-qPCR Ct value are described. For each sample, the total number of iSNVs detected is shown together with the calculation of the ratio of all iSNVs between the viral capsid region versus non-structural proteins (All iSNVs ratio) as well as for non-synonymous iSNVs only (NS iSNVs ratio). The sequencing methods used in each sample, the consensus genome coverage and the NCBI SRA accession number to the FASTQ file is shown. Individual S01-S06 were used for URT vs LRT analysis while individuals S07-S12 were used for longitudinal infection analysis. NA, not applicable; mNGS, metagenomic next-generation sequencing; PCR-tiling, amplicon-based next-generation sequencing; iSNV, intra-host single nucleotide variant.

| Individual | Sample ID | Collection date | Days since first positive sample | qRT-PCR Ct | iSNVs | All iSNVs ratio | NS iSNVs ratio | Sequencing Method | Coverage (%) | NCBI SRA Accession Number |
| --- | --- | --- | --- | --- | --- | --- | --- | --- | --- | --- |
| S01 | NW1 | 2009-08-24 | 0 | 17.5 | 0 | NA | NA | mNGS | 99.2 | SRX5167679 |
|  | BAL1 | 2009-08-24 | 0 | 22.3 | 0 | NA | NA | mNGS | 98.7 | SRX5167676 |
| S02 | NW2 | 2008-06-17 | 0 | 31.8 | 2 | NA | NA | mNGS | 100 | SRX5167677 |
|  | BAL2 | 2008-06-18 | 1 | 34.3 | 0 | NA | NA | mNGS + PCR-tiling | 97.2 | SRX5167685 |
| S03 | BAL3 | 2010-03-08 | 28 | 29.5 | 16 | 3.4 | 2.1 | PCR-tiling | 98.7 | SRX5167684 |
|  | NW3 | 2010-03-09 | 29 | 22.7 | 19 | 2.1 | NA | mNGS + PCR-tiling | 99.8 | SRX5167680 |
| S04 | NW4 | 2008-01-10 | 8 | 30 | 32 | 0.9 | 1.4 | PCR-tiling | 100 | SRX5167678 |
|  | BAL4 | 2008-01-12 | 9 | 24.1 | 18 | 1.9 | 3.1 | mNGS + PCR-tiling | 100 | SRX5167689 |
| S05 | NW5 | 2006-09-22 | 15 | 27.4 | 4 | 1.5 | NA | mNGS | 100 | SRX5167667 |
|  | BAL5 | 2006-09-27 | 20 | 23.3 | 11 | 1.3 | 1.5 | mNGS | 99.9 | SRX5167690 |
| S06 | NW6 | 2006-09-22 | 71 | 33 | 9 | 3.0 | 6.1 | mNGS + PCR-tiling | 97.8 | SRX5167668 |
|  | BAL6 | 2006-09-27 | 76 | 23 | 3 | 0.8 | 0.0 | mNGS | 99.8 | SRX5167693 |
| S07 | BAL7a | 2008-10-28 | 0 | 21.5 | 5 | 1.1 | 0.8 | mNGS+ PCR-tiling | 96.3 | SRX5167688 |
|  | NW7 | 2008-11-10 | 13 | 21.8 | 10 | 1.0 | 0.3 | mNGS | 100 | SRX5167683 |
|  | BAL7b | 2009-01-04 | 68 | NA | 19 | 1.8 | 1.5 | mNGS+ PCR-tiling | 99.9 | SRX5167687 |
| S08 | NW8 | 2008-01-10 | 0 | 20 | 3 | NA | NA | mNGS | 100 | SRX5167681 |
|  | BAL8 | 2008-01-23 | 13 | 24 | 13 | 3.4 | 12.1 | mNGS | 99.9 | SRX5167692 |
| S09 | BAL9a | 2014-05-23 | 0 | 23.5 | 10 | 2.3 | 1.0 | mNGS | 100 | SRX5167671 |
|  | BAL9b | 2014-06-04 | 12 | 26.2 | 7 | 9.0 | 4.5 | mNGS | 100 | SRX5167672 |
| S10 | BAL10a | 2009-11-03 | 0 | 21.4 | 0 | NA | NA | mNGS | 100 | SRX5167686 |
|  | BAL10b | 2010-05-12 | 190 | 23 | 19 | 2.1 | 3.8 | mNGS | 99.8 | SRX5167673 |
| S11 | BAL11a | 2010-02-19 | 30 | 20.9 | 3 | 1.5 | NA | mNGS + PCR-tiling | 97.3 | SRX5167691 |
|  | BAL11b | 2010-04-03 | 73 | NA | 18 | 1.5 | 4.1 | PCR-tiling | 97.3 | SRX5167675 |
|  | NW11 | 2010-04-13 | 83 | 24.7 | 28 | 3.8 | 25.9 | mNGS + PCR-tiling | 97.7 | SRX5167682 |
|  | BAL11c | 2010-04-20 | 90 | NA | 14 | 3.4 | 2.5 | PCR-tiling | 97.7 | SRX5167674 |
| S12 | BAL12a | 2013-01-11 | 72 | 23.8 | 12 | 1.5 | NA | mNGS | 99.7 | SRX5167669 |
|  | BAL12b | 2013-02-16 | 108 | 25.3 | 9 | 0.8 | NA | mNGS | 99.7 | SRX5167670 |

Complete and near complete HRV genomes obtained included 7 individuals with HRV-A, 3 with HRV-C, and 2 with HRV-B species (Table 1). Phylogenetic analysis showed statistically supported clusters for the consensus sequences from each individual, yielding the same HRV serotype across matched and longitudinal specimens with each individual having a different HRV serotype (Table 2, Supplemental Figure 2).

For iSNV analysis, individuals were divided into two study categories: 1) spatial niche genome evolution of URT and LRT respiratory specimens collected less than 5 days of each other (S01 –S06, Table 2), and 2) longitudinal within-host HRV genomic evolution from respiratory specimens collected across 10 to 190 days (median: 13 days) (S07 – S12, Table 2).

### No consensus genomic evolution required for HRV progression from upper to lower respiratory tract

To examine whether progression of HRV infection from URT to LRT was associated with specific HRV genomic evolution, we examined six cases where HRV-positive URT and LRT specimens were obtained within 5 days of each other, consisting of two HRV-A, two HRV-B, and two HRV-C infections (individuals S01-S06, Figure 1, Supplemental Figure 3). HRV consensus genomes had no coding changes between URT and LRT specimens for five of the six individuals. Moreover, the pair of HRV-C17 genomes (S01, Figure 1A) had no iSNVs in samples taken on the same day and the pair of HRV-C39 genomes contained only two iSNVs at 10% MAF in samples taken one day apart (S02, Supplemental Figure 3A). Individual S04 had seven consensus changes including four coding mutations in the HRV-B06 genome present in a BAL specimen taken two days after NW. All these mutations were present as iSNVs in the NW at MAFs between 20-45% (Figure 1C). Interestingly, specimens containing HRV-B genomes had significantly more iSNVs than those containing HRV-A or HRV-C sequences (ANOVA p=0.00325, Fisher-LSD test). Microbiome analysis of mNGS-sequenced specimens showed distinct microbial populations from URT and LRT specimens and a lack of oral flora present in LRT specimens, consistent with a lack of contamination between specimens that would affect URT versus LRT HRV sequence analysis (Supplemental Figure 4). Taken together, these data are consistent with the lack of requirement for specific HRV genomic evolution to allow infection of the upper and lower respiratory tract.

**Figure 1.**
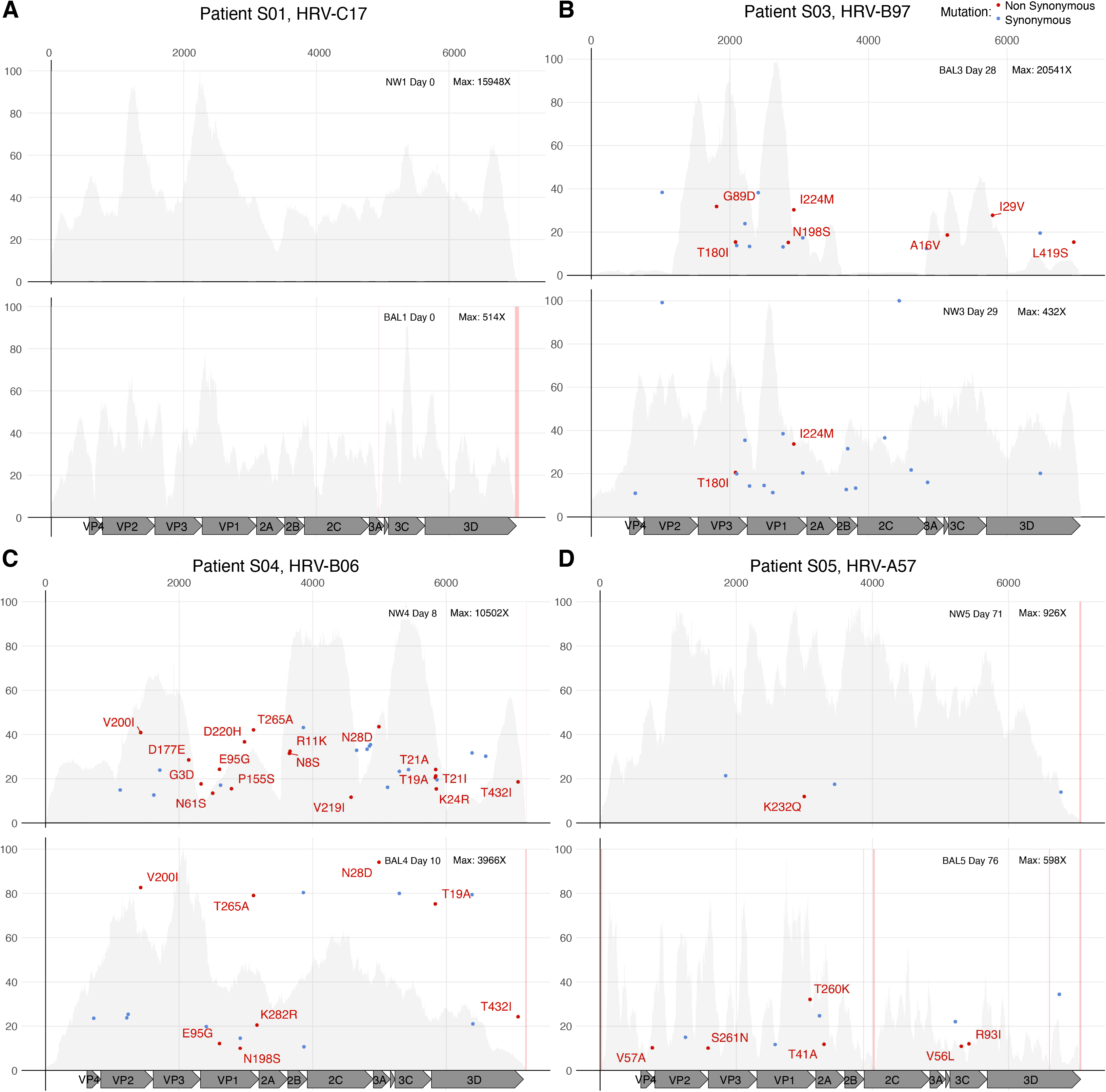
HRV evolution in upper and lower respiratory tract samples in immunocompromised individuals. Allele frequency (in percentage) of each iSNV across the HRV genome in individuals S01 (A), S03 (B), S04 (C) and S05 (D) is depicted. iSNV causing synonymous mutation are shown in blue, while non-synonymous mutations are shown in red and include the amino acid change description. Upper (NW, nasal wash) and lower (BAL, bronchioalveolar lavage) respiratory tract samples are detailed in the upper right of each plot. In addition, the day when the sample was collected relative to the first HRV positive and the maximum sequencing depth of coverage for each sample is shown. HRV mature peptides are annotated at the bottom of each panel. The grey background plot illustrates the sequencing coverage profile for each sample. Regions with low sequencing coverage (<30X) are shown in pink.

### Preferential accumulation of non-synonymous iSNVs in HRV capsid proteins

We next examined longitudinal evolution of HRV genomes across six individuals (5 HRV-A and 1 HRV-C; individuals S07-S12 in Table 2). Given the lack of mutational changes present in closely matched URT and LRT specimens above, we utilized both URT and LRT specimens for this analysis. Samples were collected a median of 30 days after the first detection of HRV in each individual (range 0-190 days). iSNVs present in the first collected specimen increased in allele frequency to near fixation in five of the six individuals (Figure 2, Supplemental Figure 5), with the sole exception being individual S10 who had no iSNVs in the day 0 sample and only a day 190 specimen available.

**Figure 2.**
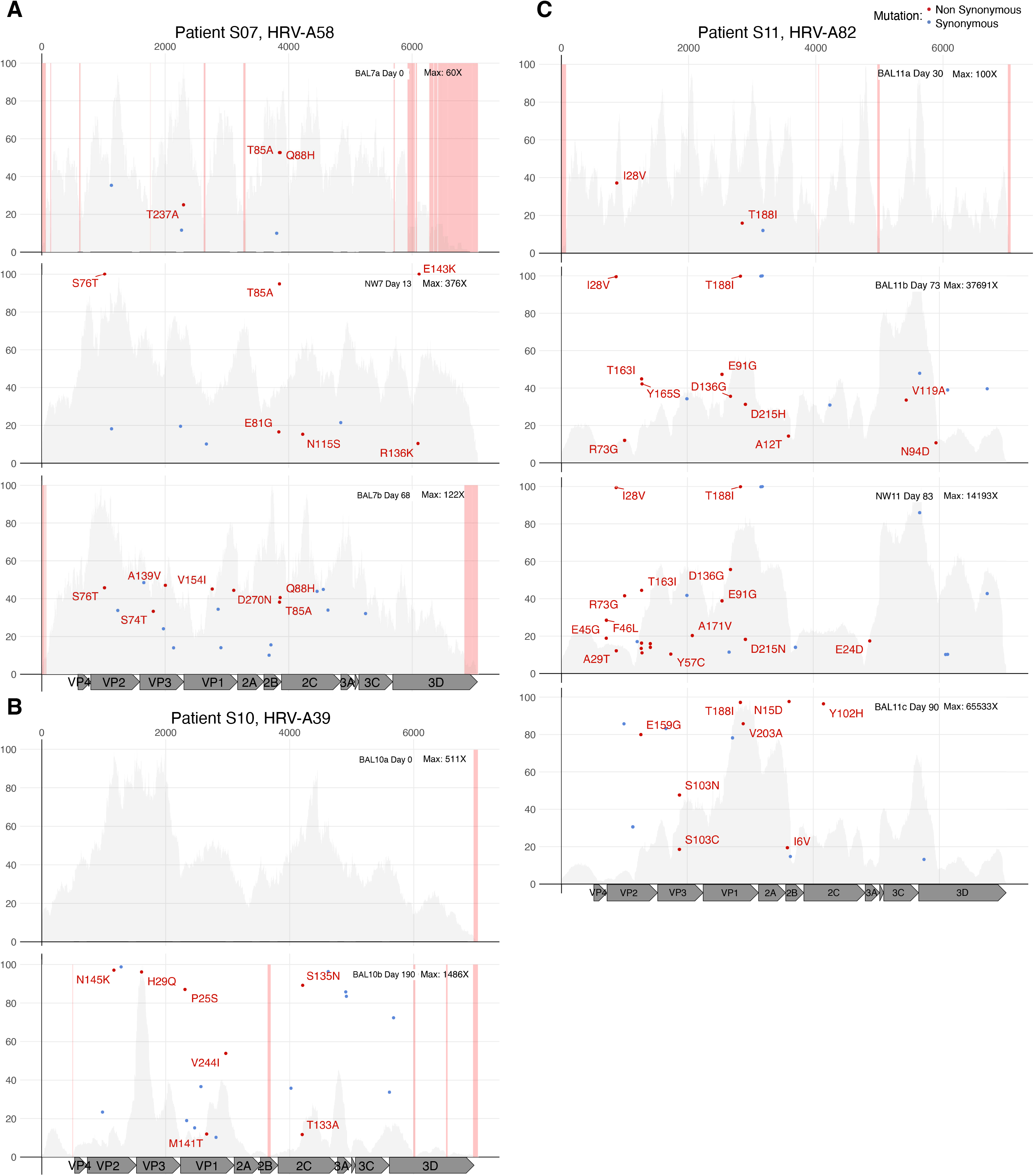
Long-term HRV infection dynamics in immunocompromised individuals. Minor allele frequency (in percentage) of each iSNV across the HRV genome in individuals S07 (A), S10 (B) and S11 (C) is represented by dots. Blue dots indicate iSNV causing synonymous mutation and red dots indicate non-synonymous mutation which also includes the amino acid change description. Upper (NW, nasal wash) and lower (BAL, bronchioalveolar lavage) respiratory tract samples are detailed in the upper right of each plot. In addition, the day when the sample was collected relative to the first HRV positive and the maximum sequencing depth of coverage for each sample is shown. HRV mature peptides are annotated at the bottom of each panel. The grey background plot illustrates the sequencing coverage profile for each sample. Regions with low sequencing coverage (<30X) are shown in pink.

Examination of these six longitudinal infections revealed a significant accumulation of non-synonymous iSNVs in HRV capsid proteins compared to non-structural proteins (Supplemental Figure 6). Across the six infections, a total of 65 non-synonymous iSNVs in capsid proteins were detected compared to 23 in non-structural proteins. When iSNVs from all twelve individuals were included, the median capsid-to-nonstructural ratio for nonsynonymous iSNVs was 2.3 (range: 0 -25.9) and for total iSNVs was 1.8 (range: 0.8 – 9.0), after adjusting for locus length (Table 2).

Given the high proportion of HRV-A samples in our study, we specifically modeled iSNVs located in the HRV-A16 capsid structure (PDB: 1AYM) over three time periods relative to the first HRV-A positive sample: day 0, 12-30 days, and >68 days (Figure 3). Capsid iSNVs accumulated in both number and allele frequency across time, appearing first in VP2 and VP3 proteins and later in VP1 (Figure 3). Among HRV-As, surface resides on the capsid were 4.0 times more likely to contain iSNVs compared to non-surface residues (p=4.7e-6, Pearson’s chi-squared test), and surface residues were 3.1 times more likely to contain iSNVs across HRV-A/B/C (p=1.3e-5). Allele frequency of capsid iSNVs showed a non-significant positive association with residue solvent accessibility surface area for VP1 and VP3, but not for VP2 (Supplemental Figure 7).

**Figure 3.**
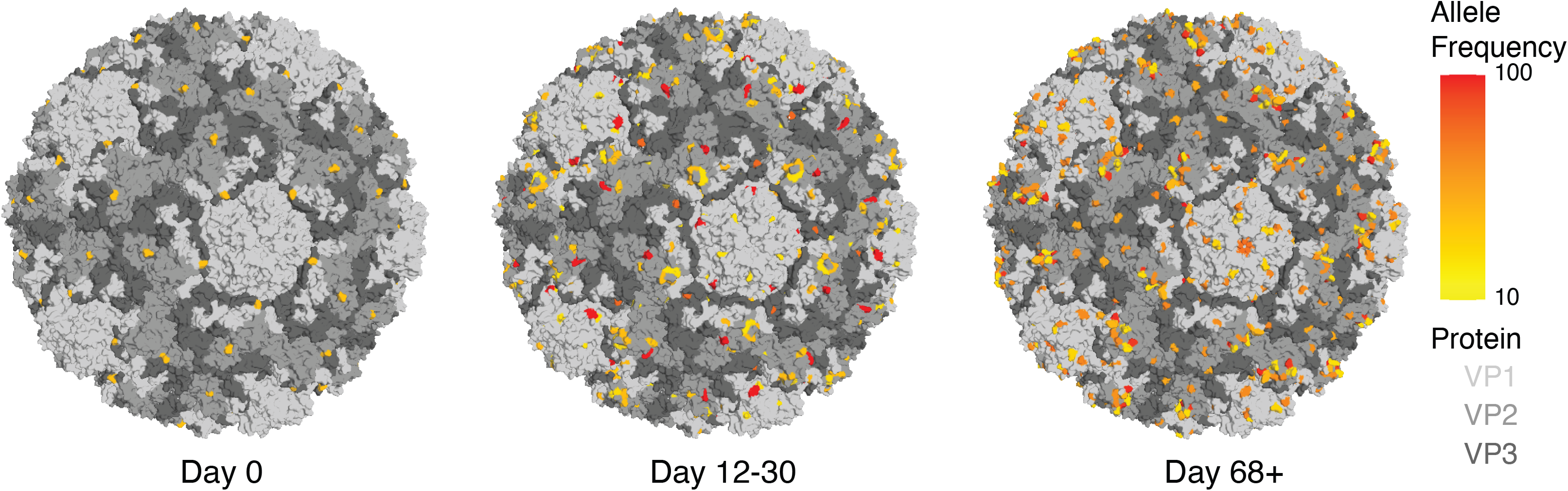
HRV-A capsid evolution dynamics during prolonged within-host infection. iSNVs modeled in the 3D crystal structure of the HRV-A16 capsid (PDB accession number 1AYM). Proteins VP1, VP2 and VP3 are differentiated in the 3D model with different gray degrees. iSNV allele frequency is depicted on a scale from yellow to red (from 10% to 100%). iSNV detection was categorized in day zero (first sample positive detected), after 12 to 30 days from the first HRV positive and after more than 68 days from the first HRV positive.

### iSNVs are enriched in sites of higher in vitro mutational fitness effect in VP2 and not located in HRV epitope sites

To understand potential causes or effects of emergent iSNVs, we compared iSNV presence with estimations of mutational fitness effect (MFE) from deep mutational scanning of the Coxsackievirus B3 capsid [30]. iSNVs sites from all HRV species had a higher average MFE per residue VP2 (p-value = 0.004) but not VP1, VP3, and VP4 (p-value > 0.05, Mann Whitney Wilcoxon) (Supplemental Figure 8). iSNVs were not preferentially located in neutralizing antibody epitopes previously mapped in HRV-A2 and HRV-B14 (p=0.192, Chi-square test) [31,32].

### Newly increasing non-synonymous iSNV haplotypes detected in future consensus HRV sequences

We next examined whether iSNVs that increased in allele frequency in our 2006-2014 specimens could be detected in other HRV consensus sequences present in NCBI GenBank. Sequence haplotypes that nearly fixed in two longitudinal HRV-A infections from 2006-2014 were found in NCBI GenBank sequences identified in future years (Supplemental Figure 9). For example, the HRV-A58 haplotype (VP2: S76T, 2B:T85A, 3D:E143K) that nearly fixed in individual S07 from November 2008 was detected in five HRV-A58 sequences from Seattle in 2021 as well as one pediatric infection from Wisconsin in 2009 [33]. Similarly, the HRV-A82 haplotype (VP2:I28V, VP1:T188I) that fixed in individual S11 in April 2010 was also found in HRV-A82 sequences taken from 2009 and 2010 in Wisconsin and 2016 and 2020 from Seattle [33,34]. Though iSNV VP2:N145K from S10 could also be detected in a 2021 HRV-A39 sequence taken from Seattle, it did not have a close phylogenetic relationship with BAL9b and failed to contain the full iSNV haplotype.

## Discussion

Here, we compared the within-host genomic evolution of HRV in URT and LRT specimens from 12 immunocompromised individuals after HCT. Overall, we found a lack of specific genetic changes associated with compartmentalization in URT versus LRT and a strong genetic plasticity in capsid proteins compared to non-structural proteins. Capsid iSNVs were preferentially located in surface residues as opposed to core residues, but intriguingly these residues were not necessarily associated with neutralizing antibody epitopes or had particularly strong fitness effects in deep mutational scanning data from CVB3 capsid.

Global epidemiology of HRV has shown that HRV-A is the most frequently detected species in both immunocompetent and immunocompromised individuals [14,17,35,36], matching our preferential recovery of HRV-A in more than half of the infections profiled. In addition, no prevalence of a specific HRV serotype was found, albeit in only 12 infections.

Intra-host evolution of HRV infections has been less studied compared to infections with enveloped human respiratory viruses. Cordey et al. (2010) examined intra-host evolution after experimental infection of HRV-A39 over 5 days in fewer than five immunocompetent adults [37]. They found most low-level mutations occurred in VP2, VP1, and 2C regions and most consensus level mutations occurred in VP1, VP2, and VP3 structural proteins. Tapparel and colleagues (2011) also found that structural proteins underwent a higher mutation rate during chronic HRV-A and HRV-B infections in five lung transplant recipients [38]. In addition, they found no ecological association between HRV species and detection in upper and lower respiratory tract specimens. Our results match these studies in finding preferential accumulation of mutations in the capsid region compared to non-structural region in longitudinal infections. Our results also support the lack of association of LRT infection with any specific mutation or genotype, using HRV sequencing data from matched LRT and URT combined with metagenomic analysis to ensure specimen integrity. Unlike Cordey et al., we detected mutations in the capsid drug-binding pocket – a hydrophobic cavity in the capsid conferring antiviral drug susceptibility – despite the patients having not received any antiviral treatment [39]. Specifically, for both individuals with HRV-B, N198S in VP1 protein was detected. In addition, individual S04 had the mutation D200H in VP1 and individual S03 had the mutation I224M at VP1, which is a confirmed residue that interacts with an antiviral capsid-inhibitor called pleconaril [12]. A mutation in the drug binding-pocket was also detected in an HRV-A57 sample of individual S05.

The lack of a specific evolutionary requirement for HRV to change LRT/URT niche strongly indicates HRV progression to LRTI is largely due to non-viral factors, which is consistent with prior associations of LRT progression with intravenous immunoglobulin administration, steroid use, and low monocyte count [40]. While LRT samples could reflect URT secretions that were aspirated during the sampling procedure, we evaluated this bias by assessing a comparative microbiome analysis in the NW and BAL with metagenomic NGS. Microbiome profiles in BAL at advanced infection times showed high abundance of both commensal and opportunistic pathogens in the immunocompromised host (*Stutzerimonas stutzeri*, *Brevundimonas*, *Alcaligenes*, *Ralstonia*, and *Rothia mucilaginosa*) [41–44].

Though limited to infections in only two individuals, our sequencing data supported a higher intra-host diversity of HRV-B infections. Intriguingly, multiple studies have found HRV-B infections to have lower viral loads and/or reduced symptom severity compared to HRV-A or HRV-C infections [45–47]. Our HRV-B infections had higher Ct values compared to HRV-A or HRV-C infections (Table 2), which may result in higher allele frequencies in mixed infections [48]. Alternatively, the lower severity of HRV-B infections may bias or complicate ascertainment of the initiation of infection and potentially lead to longer infections that allow greater intra-host diversity. Given the reduced frequency of HRV-B infections, comparatively less effort has gone into understanding evolutionary rates and factors of HRV-B and more work is required to understand inter-species differences in rhinovirus evolution.

We also examined whether newly derived capsid mutations and sequencing haplotypes could be found in other HRV genomes based on searches of NCBI GenBank. Many of these HRV consensus genomes in NCBI GenBank are derived from our laboratory from 2021-22 after screening the high number of COVID-19-negative nasal swab specimens sent to our lab and sequencing the HRV positives. Indeed, several newly-derived HRV mutations from our study patients could be found in future community sequences. While on the surface this matches results seen with sequencing longitudinal influenza A virus infections in immunocompromised individuals [49], given the lack of antigen drift seen in rhinoviruses [50], these data could just be indicative of the limited number of sequences available and viable mutations for the capsid of a given HRV serotype. Indeed, new intra-host mutations arising in one our profiled individuals from 2010 were also seen in HRV from Wisconsin in 2009/2010. Certainly, the amount of sequencing data on HRV pales in comparison to that available for influenza virus, complicating confident determination of global allele frequencies of HRVs, especially after dividing across all HRV serotypes.

Our examination of the association between iSNV presence and mutational fitness effect in CVB3 deep mutational scanning data only recovered a modest relationship with iSNVs present in the VP2 protein and failed to find an association with iSNV allele frequency [30]. Although these results suggest an important role of VP2 in the HRV-A intra-host evolution, *in vitro* experiments are needed to confirm this hypothesis. The lack of association between mutational fitness effect and iSNV presence in other capsid proteins as well as the lack of association with iSNV allele frequency may be the result of competition between equally fit viral populations in an environment with a lower selection pressure or simply due to the inability to accurately model and relate residues across different enterovirus species.

Our study was chiefly limited by the number of individuals and specimens examined. Individuals were not regularly sampled over space and time and not all remnant specimens were stored from the clinical care. Given the incredible diversity of HRV serotypes, our limited sampling constrains our ability to make generalizable conclusions about HRV evolution. In addition, capsid structures and deep mutational scanning data are only available for select serotypes or species [29,30]. Although the first HRV-positive sample was considered representative of the beginning of the infection, it is not possible to confirm this with certainty. However, clinical practice guidelines recommend patients report symptoms urgently and then undergo PCR testing. We also did not perform two independent library preparations for each specimen sequenced, though we controlled for this by demonstrating robustness of our library preparation and sequencing protocols for select specimens and using a relatively high iSNV allele frequency of 10% [48].

Overall, our work provides valuable insights into the within-host HRV evolution including the higher ratio of variability of the capsid proteins compared to non-structural proteins, the mutation-independent compartmentalization of URT and LRT infections, and the identification of emerging within-host mutations in the immunocompetent community. Our data highlight the unique evolutionary constraints seen in HRVs – and their seemingly nihilistic evolution – that are generally not present in infections with enveloped human respiratory viruses.

## Data availability

Sequencing data obtained in this study is available at GenBank in NCBI BioProject PRJNA907865.

## Conflict of interest

ALG reports contract testing from Abbott, Cepheid, Novavax, Pfizer, Janssen and Hologic and research support from Gilead and Merck, outside of the described work. AW reports research support from Pfizer, Amazon, GlaxoSmithKline/Vir, Ansun Biopharma, and Allovir; receiving personal fees from Kyorin Pharmaceutical and Vir Biotechnology; and receiving grants from Amazon and VB Tech, all outside the submitted work.

## Funding

This work was supported by the Department of Laboratory Medicine and Pathology of the University of Washington and NIH grant K23 AI114844-02 (A.W.).

## Supplemental Table Legends

**Supplemental Table 1. All iSNVs detected across 12 HRV infections in immunocompromised individuals**

For each individual the intra-host single nucleotide variation (iSNV) is described. For each iSNV the position in the rhinovirus genome, the protein affected, the minor allele frequency and the amino acid change (if appropriate) is detailed in a spreadsheet per sample. The NCBI SRA accession number to the fastq file for each sample analyzed is detailed in the main header. BAL: bronchoalveolar lavage. NW: nasal wash.

**Supplemental Table 2. Deduplicated coding iSNVs as mapped to HRV-16 capsid structure (PDB: 1AYM).** Unique capsid iSNV positions from HRV-A, B, and C infections along with their corresponding position and structure metadata based on in HRV-16 capsid (PDB: 1AYM) are listed, along with other sequencing metadata and neutralizing epitope residues used for statistical tests.

## Supplemental Figure Legends

**Supplemental Figure 1. Comparison of iSNV detection by amplicon-based and metagenomic sequencing.** Scatter plots show the allele frequency (in percentage) of a given iSNV found in sample BAL11b during different rounds of amplicon-based sequencing (A) and among both amplicon-based and metagenomic sequencing for samples NW11 (B), BAL4 (C) and NW3 (D).

**Supplemental Figure 2. Phylogenetic classification of HRV samples.** Maximum likelihood inference of the VP1 nucleotide sequence including reference sequences of all HRV-A, HRV-B and HRV-C serotypes. The name of the consensus sequences for HRV from each sample is highlighted in red. The HRV serotype in each reference is informed in the tip name. Statistical support of relevant phylogenetic clades’ association (SH-alrt / UF-Bootstrap) are indicated. HRV-A, HRV-B and HRV-C clades are denoted with the branch colors.

**Supplemental Figure 3. HRV evolution in upper and lower respiratory tract samples comparison at short infection times in immunocompromised individuals.** Minor allele frequency (in percentage) of each iSNV across the HRV genome in individuals S02 (A) and S06 (B) is represented by dots. Blue dots indicate iSNV causing synonymous mutation and red dots indicate non-synonymous mutation which also includes the amino acid change description. Upper (NW, nasal wash) and lower (BAL, bronchioalveolar lavage) respiratory tract samples are detailed in the upper right of each plot. In addition, the day when the sample was collected regarding to the first HRV positive and the maximum sequencing depth of coverage for each sample is informed. HRV secondary proteins annotation are indicated at the bottom of the graphics. Grey plot at the background informs the sequencing coverage profile for each sample. Pink-colored regions denotes low sequencing coverage, thereby not analyzed.

**Supplemental Figure 4. Microbiome characterization in upper and lower respiratory tract samples from immunocompromised individuals during HRV infection.** A) NMDS ordination with Bray-Curtis dissimilarity matrix based on the relative abundance of bacteria genus communities. Ellipse of 95% confidence level for the multivariate t-distribution were constructed by using the ’stat_ellipse’ function of ggplot2 package to illustrate the lower (BAL) respiratory tract samples. No 95% ellipse could be estimated for upper (NW) respiratory tract samples due to the low number of samples. B) Bar-plots show the relative abundance of bacterial genus found in each sample.

**Supplemental Figure 5. Long-term HRV infection dynamics in immunocompromised individuals.** Minor allele frequency (in percentage) of each iSNV across the HRV genome in individuals S08 (A), S09 (B) and S12 (C) is represented by dots. Blue dots indicate iSNV causing synonymous mutation and red dots indicate non-synonymous mutation which also includes the amino acid change description. Upper (NW, nasal wash) and lower (BAL, bronchioalveolar lavage) respiratory tract samples are detailed in the upper right of each plot. In addition, the day when the sample was collected regarding to the first HRV positive and the maximum sequencing depth of coverage for each sample is informed. HRV secondary proteins annotation are indicated at the bottom of the graphics. Grey plot at the background informs the sequencing coverage profile for each sample. Pink-colored regions denotes low sequencing coverage, thereby not analyzed.

**Supplemental Figure 6. Non-synonymous and synonymous iSNVs percentage in the capsid and the non-structural proteins.** The percentage of sites in the capsid proteins (positive values) and the non-structural proteins (negative values) where an iSNV was detected per sample per day of collection is shown, for both iSNVs causing a non-synonymous (A) and synonymous (B) mutation. Samples are colored by individual including a 50% transparency effect for the non-structural protein results.

**Supplemental Figure 7. Allele frequency for each iSNV compared to solvent accessibility surface area for the corresponding VP1, VP2 and VP3 residue in HRV-A16 capsid (PDB: 1AYM).** Comparison of minor allele frequency (AF) in percentage and the solvent accessibility surface area (SASA) in percentage for each iSNV located in each of the capsid proteins of HRV-A samples. Colors in dots differentiate the time categorization used in Figure 3. Regression lines are shown for each protein, together with the estimated equation and the square ratio value.

**Supplemental Figure 8. Evaluation of the mutational fitness effect in a CVB3 capsid (PDB: 4GB3) HRV-A, B, and C iSNVs.** Violin plot of the average mutational fitness effect (MFE) in sites with detection of non-synonymous iSNVs in the VP1, VP2, VP3 and VP4 capsid proteins across all 12 HRV infections. P-value of the ranked Wilcoxon test is displayed.

**Supplemental Figure 9. Maximum likelihood inference of HRV-A complete genome alignment including all sequences published in NCBI GenBank up to January 2023.** The HRV serotypes in which classified the studied sequences are informed. Color in tip names denotes the year of collection of the sample. Sequences from the analyzed individuals are indicated with an arrow. Phylogenetic clades of serotypes containing sequences sharing the high-frequency haplotype of each individual’s samples are zoomed, and those sequences are denoted with a black circle.

## Notes

https://github.com/greninger-lab/HRV_Longitudinal

