## Supplemental material for "Within-host rhinovirus evolution in upper and lower respiratory tract highlights capsid variability and mutation-independent compartmentalization"

**Supplemental Methods**

**Metagenomic and Amplicon-based Sequencing**

Metagenomic next-generation sequencing (mNGS) libraries were prepared as described previously using RNA extracted from 50µL of NW and BAL with Zymo Viral RNA kit (Zymo Research) [1,2]. Briefly, extracted RNAs were treated with Turbo DNase I (Thermo Fisher) and cDNA was generated using SSIII Reverse Transcriptase (Thermo Fisher). Then, double stranded DNA was generated using Sequenase V2.0 DNA Polymerase (Thermo Fisher) and purified using DNA Clean & Concentrator (Zymo Research). mNGS libraries were performed using two-fifths volumes of Nextera XT DNA Library Preparation Kit (Illumina). Libraries were run on 1x192 bp cartridge on Illumina MiSeq.

Samples without optimal mNGS performance were completed by PCR-tiling method using specific primers sets designed from the consensus genome of the paired sample. Briefly, overlapped amplicons of ~1kb length covering the whole genome were performed using the Qiagen One-Step RT-PCR with the following cycle conditions: reverse-transcription incubation of 5 min at 37°C, 30 min at 55°C and PCR amplification of 15 min at 95°C followed by 40 cycles of PCR with 30 sec at 94°C, 30 sec at 55°C and 90 sec at 72°C and with last elongation step of 2min at 72°C (Appendix Table 1). The amplicons were quantitated and pooled equimolarly per sample, then cleaned up with 0.8X AMPure XP (Beckman Coulter). Libraries were performed using Nextera XT DNA Library Preparation Kit (Illumina) and run on 2x300 bp and 1x192 bp cartridges on an Illumina MiSeq.

**Sequencing Data Analysis**

Sequencing FASTQ files were adapter- and quality- trimmed with Cutadapt v4.2 [3] and mapped against a concatenation of all existing rhinovirus complete genomes informed in ICTV using Revica pipeline (<https://github.com/greninger-lab/revica>) to obtained the consensus genome of each sample [4]. In addition, based on the consensus genome of the first positive sample detected per patient, the presence and frequency of intra-host single-nucleotide variant (iSNVs) was analyzed using LAVA pipeline [5]. iSNV was considered with a minimum frequency of 10% in the positions with a minimum sequence depth of coverage of 30X. Mutations were annotated according to secondary protein annotations of NC_038311.1 for rhinovirus A1, NC_001490.1 for rhinovirus B14 and NC_009996.1 for rhinovirus C. All sequencing reads files analyzed are available at NCBI SRA database as well as the majority consensus in NCBI GenBank (Appendix Table 2).

**HRV phylogenetic analysis**

HRV serotyping was determined phylogenetically with reference sequences downloaded from NCBI GenBank according to the ICTV classification [4]. Alignments of the VP1 region were performed including the consensus sequences of each analyzed sample to assign the HRV serotype. Additionally, complete genomes alignment by HRV specie including all HRV published in NCBI GenBank up to January 2023 were performed to assess the relationship of the studied samples with contemporaneous and future HRVs. All the maximum likelihood trees were inferred using IQ-TREE v2.1 [6]. The molecular evolution model was estimated with ModelFinder and the reliability of sequences’ clusters was evaluated using SH-aLRT test (1000 replicates) and UFBoot2 method (10000 replicates) [7–9]. Monophyletic clusters were considered when SH-aLRT value ≥80% and UFbootstrap value ≥ 90%. Phylogenetic tree visualization was performed with Figtree v1.4.4 (<https://github.com/rambaut/figtree/releases>). To evaluate if any of amino acid changes that emerged and fixed during within-host evolution was also present among HRV from the community, the region of the open reading frame containing the iSNVs was translated and tblastn function in NCBI was used to search among rhinovirus sequences (taxid: 12059) from translated nucleotide databases those including the mutations of interest.

**Protein modeling analysis**

To further analyze the iSNVs causing nonsynonymous changes in the VP proteins, the amino acid residue for each iSNV was localized on the HRV-A16 capsid structure (PDB: 1AYM) using HHPred and visualized using Pymol [10,11] (Annote_Pymol_Position.py). iSNVs were highlighted on the capsid crystal structure based on the minor allele frequency (Add_Colors_Pymol.py). The function ‘surface accessibly per residue’ in Pymol was used to calculate the solvent accessibility surface area (SASA) for each residue containing an iSNV was detected. Surface versus non-surface capsid residues were also determined by ViperDB v3.0 analysis [12].

To associate iSNVs with in vitro mutational fitness effect (MFE) data, we used a recent deep mutational scan of the Coxsackievirus B3 capsid [13,14]. HRV-A16 (PDB: A1YM)-based loci for the HRV-A iSNVs along with HRV-B and HRV-C iSNVs were localized on the Coxsackievirus B3 capsid (PDB: 4GB3) using HHPred (DeepMutation_Script.py) [14]. iSNVs that localized to alignment gaps were removed for the analysis. Average MFE per residue was calculated using all three replicate experiments and a two-sided Mann–Whitney statistical test was used to test for association with iSNVs.

**Microbiome analysis**

Microbiomes from samples underwent mNGS-only were analyzed to rule out cross-contamination between BAL and NW during sampling. Quality-filtered FASTQ files were analyzed with Kaken2 using the standard prebuilt database (v20221209) [15]. Kraken2 reports were combined in an out table using kraken-biom, which was then filtered according to Bacteria kingdom. OTU matrix was normalized by rarefaction and Bray-Curtis dissimilarity based on NMDS method and the taxonomy relative abundance per sample were calculated with phyloseq library in RStudio [16,17]. Graphics were performed with ggplot2 library [18].

**Appendix Table 1. Primer sequences for HRV genome amplification.** Sequences designed to amplify and sequence genomic regions with low coverage from the metagenomic methodology.

| Primer Name | Primer sequence (5' - 3') |
| --- | --- |
| HRV34_1F | TGCCAGTTTTATTTCCCTCCC |
| HRV34_1R | GTGTTCTGGTATCATAGCAAC |
| HRV34-2F | AAGCAAATTCCATCAAGGTACTT |
| HRV34-2R | CATGTGTAAATTCGTGTCACG |
| HRV34-3F | GGGTGCAAGGATTTTTGTCTAA |
| HRV34-3R | TCATGGCTTGTGACTTTAATTGG |
| HRV34-4F | GAAGCAACTTATTACTGTAGACA |
| HRV34-4R | CCTTATCTGGTAAATCTGCCAT |
| HRV34-5F | TGTCAGATGGTGTCAAGTGTT |
| HRV34-5R | TGCTAGGTAGACCACATTTATCT |
| HRV34-6F | GTTTTGCTGCCATGTTACTAAGA |
| HRV34-6R | AGAAGTAAGTCATAAGGAGGGA |
| HRV9-1F | TTAAAACAGCGGATGGGTATC |
| HRV9-1R | AAGAAGGCAGCCACTATGGAA |
| HRV9-2F | GCTCAAAAGGCTGGTGTTG |
| HRV9-2R | ACTGAAGCTAATGTCTGTTTCGT |
| HRV9-3F | ACAAACTGATGCCCTGACAGA |
| HRV9-3R | CCTCCTGCTGTTAACAAACCT |
| HRV9-4F | AGGCGATTGTGGTGGTATCT |
| HRV9-4R | TGACATTGTTGCGTTAAGCTTTC |
| HRV9-5F | GACATCTGTTTGCACAGTACTTA |
| HRV9-5R | CAGCATGTTTTCAGTGGGCT |
| HRV9-6F | AATGTCTTCCCGGGTAGC |
| HRV9-6R | CTTCTATTTCTAATCTAAACTAGAGG |
| HRV-44midF | CACATGACAACAAGGACACAG |
| HRV-44midR | TCCATAGACTGTGTCCTTGTT |
| HRV44-3-2-nestedF | TGGGTAGCATAGCCTTTAGAG |
| HRV44-3-2-nestedR | AACCTGTTCCAAATGCCTTCC |
| HRV40-1F | CTTACGGGAGTTGTACTCTGTTA |
| HRV40-1R | ACATTCCCATATGTTGCGCTAG |
| HRV40-2F | CTTGGCAGGAGTGGTTACA |
| HRV40-2R | CATCCAACGCAGGTGCTGA |
| HRV40-3F | CTGTGCTTTGTGTCAGGATGT |
| HRV40-3R | CACAGTGAAAGTGTCTAAGATCA |
| HRV40-4F | CTGGTGATTGTGGAGGTAAATT |
| HRV40-4R | AAGACGCAGGCAAGTCATAG |
| HRV40-5F | CCCGTTTATATGTGGTAAAGCAG |
| HRV40-5R | GTAACCATAGGTAAGTCAACAC |
| HRV40-6F | CCTTATGTTGCTATGGGGATC |
| HRV40-6R | CATACCACTCGTGCAGGAGT |
| HRV40-6bisR | TCATAAGGAGGGATATACAGTG |
| HRV49-1F | TATCGCAACTTAGAAGCTTGAG |
| HRV49-1R | ATGCTGTTATGGTGTGGCCT |
| HRV49-2F | GGTCAGAACATGTACTATCATTC |
| HRV49-2R | AAGAGCTGATCGGCCGAGA |
| HRV49-3F | AGACACTCAAGCCTCAGGA |
| HRV49-3R | CTAATAAGGCGAGGGTAGC |
| HRV49-4F | GCACTTCCATTACTGATAAGC |
| HRV49-4R | GGCCATTCTCTTCACAGTATTTA |
| HRV49-5F | CGCAAATGAGAGAAGAGCTC |
| HRV49-5R | ATGGGAAGTATCATTAATGCTGC |
| HRV49-6F | CCACCAGAGATGTGAGCAA |
| HRV49-6R | CCTACCCACTGATGTAGTTCT |

**Appendix table 2. NCBI accession numbers and metadata.** For each individual the NCBI GenBank accession number to the majority consensus in FASTA format and the NCBI SRA raw sequence data in FASTQ is detailed. For each case the collection date of the sample, serotype and sequencing method (mNGS, metagenomic NGS) is described.

| Patient | Sample ID | Collection date (yyyy-mm-dd) | Serotype | Sequencing Method | FASTA Accession Number | FASTQ Accession Number |
| --- | --- | --- | --- | --- | --- | --- |
| S01 | NW1 | 2009-08-24 | C17 | mNGS | MZ667421 | SRX5167679 |
|  | BAL1 | 2009-08-24 |  | mNGS | OQ536576 | SRX5167676 |
| S02 | NW2 | 2008-06-17 | C36 | mNGS | MZ438010 | SRX5167677 |
|  | BAL2 | 2008-06-18 |  | mNGS +PCR-tiling | OQ536575 | SRX5167685 |
| S03 | BAL3 | 2010-03-08 | B97 | PCR-tiling | MZ667418 | SRX5167684 |
|  | NW3 | 2010-03-09 |  | mNGS + PCR-tiling | OQ536578 | SRX5167680 |
| S04 | NW4 | 2008-01-10 | B06 | PCR-tiling | MZ667416 | SRX5167678 |
|  | BAL4 | 2008-01-12 |  | mNGS + PCR-tiling | OQ536573 | SRX5167689 |
| S05 | NW5 | 2006-09-22 | A57 | mNGS | MZ667415 | SRX5167667 |
|  | BAL5 | 2006-09-27 |  | mNGS | OQ536572 | SRX5167690 |
| S06 | NW6 | 2006-09-22 | A105 | mNGS + PCR-tiling | MZ542285 | SRX5167668 |
|  | BAL6 | 2006-09-27 |  | mNGS | OQ536571 | SRX5167693 |
| S07 | BAL7a | 2008-10-28 | A58 | mNGS+ PCR-tiling | OQ536569 | SRX5167688 |
|  | NW7 | 2008-11-10 |  | mNGS | OQ446464 | SRX5167683 |
|  | BAL7b | 2009-01-04 |  | mNGS+ PCR-tiling | OQ536570 | SRX5167687 |
| S08 | NW8 | 2008-01-10 | A78 | mNGS | MZ667417 | SRX5167681 |
|  | BAL8 | 2008-01-23 |  | mNGS | OQ536574 | SRX5167692 |
| S09 | BAL9a | 2014-05-23 | A102 | mNGS | MZ458533 | SRX5167671 |
|  | BAL9b | 2014-06-04 |  | mNGS | OQ536566 | SRX5167672 |
| S10 | BAL10a | 2009-11-03 | A39 | mNGS | MZ667419 | SRX5167686 |
|  | BAL10b | 2010-05-12 |  | mNGS | OQ446463 | SRX5167673 |
| S11 | BAL11a | 2010-02-19 | A82 | mNGS + PCR-tiling | MZ667414 | SRX5167691 |
|  | BAL11b | 2010-04-03 |  | PCR-tiling | OQ536567 | SRX5167675 |
|  | NW11 | 2010-04-13 |  | mNGS + PCR-tiling | OQ536577 | SRX5167682 |
|  | BAL11c | 2010-04-20 |  | PCR-tiling | OQ536568 | SRX5167674 |
| S12 | BAL12a | 2013-01-11 | C28 | mNGS | MZ447875 | SRX5167669 |
|  | BAL12b | 2013-02-16 |  | mNGS | OQ536565 | SRX5167670 |
