## Supplementary figures and images for "Within-host rhinovirus evolution in upper and lower respiratory tract highlights capsid variability and mutation-independent compartmentalization"

### Supplemental figure 1

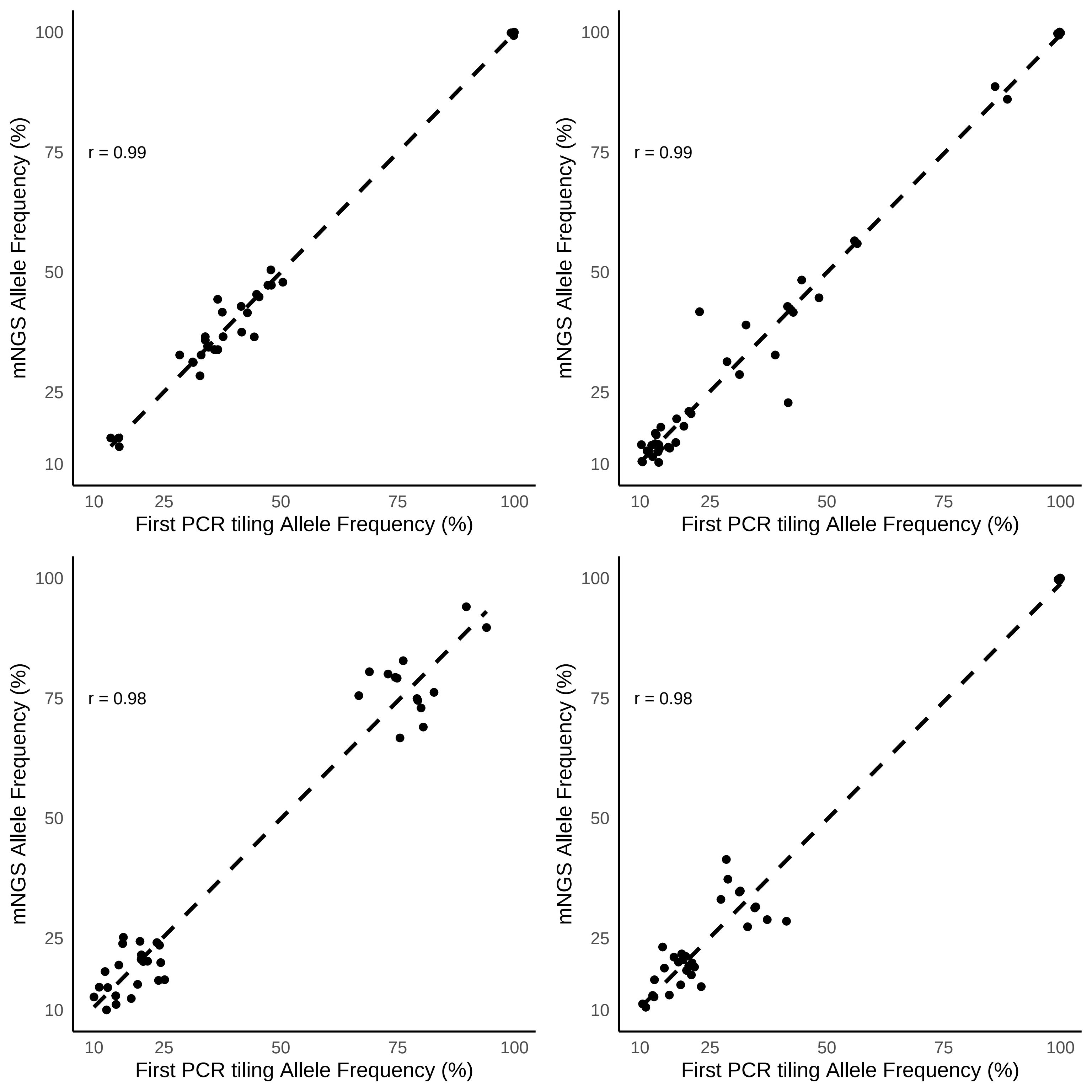

### Supplemental figure 2

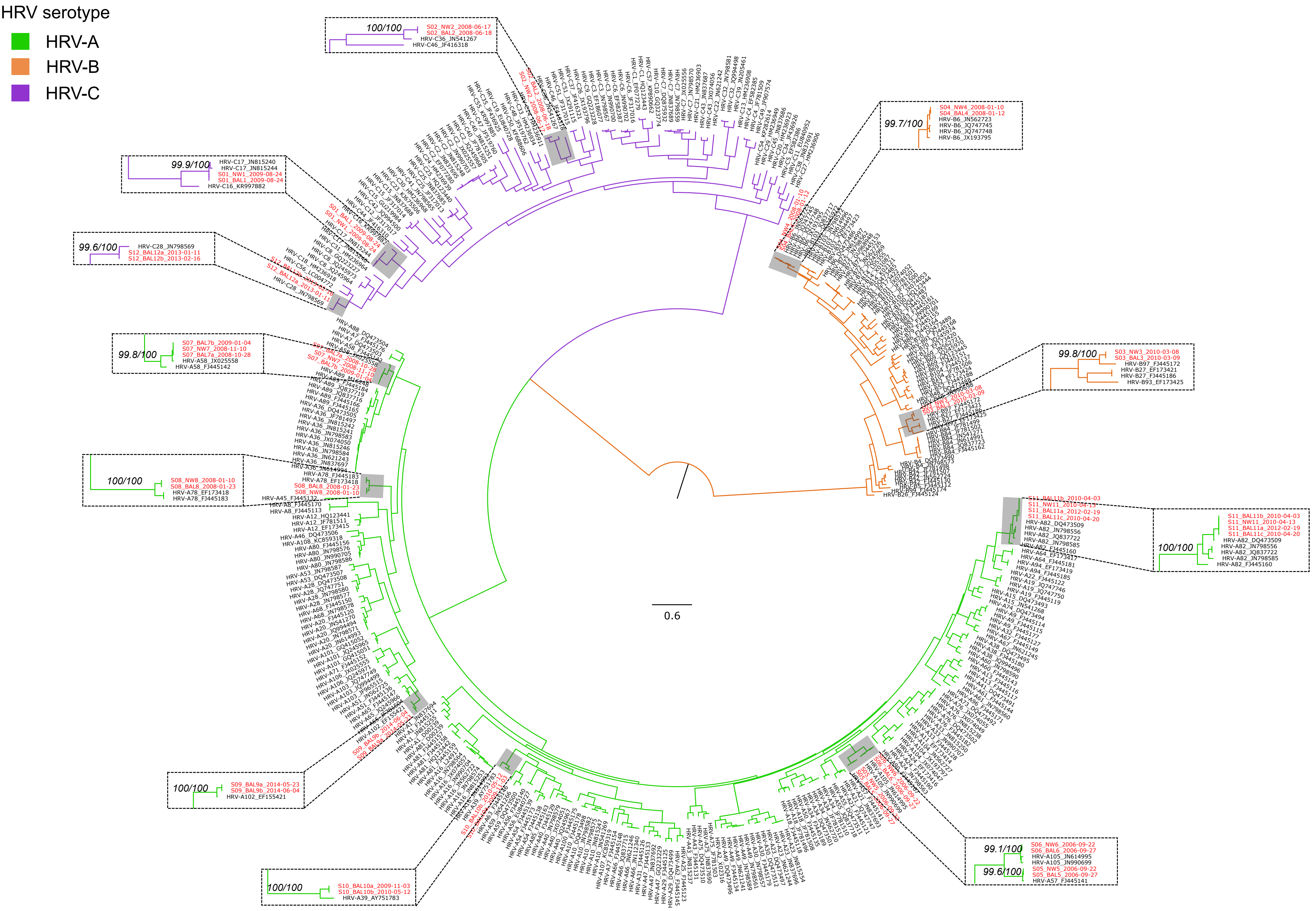

### Supplemental figure 3

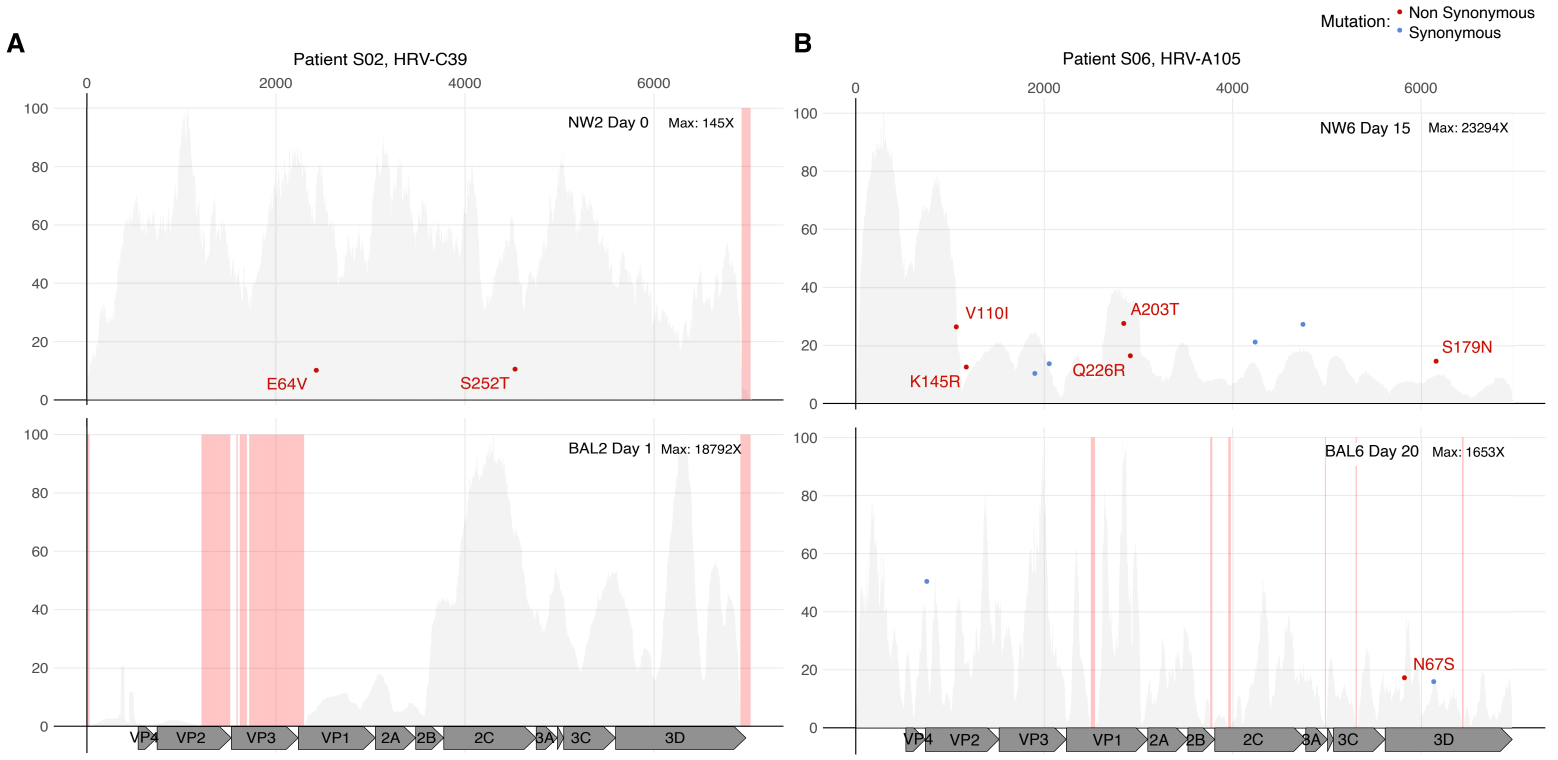

### Supplemental figure 4

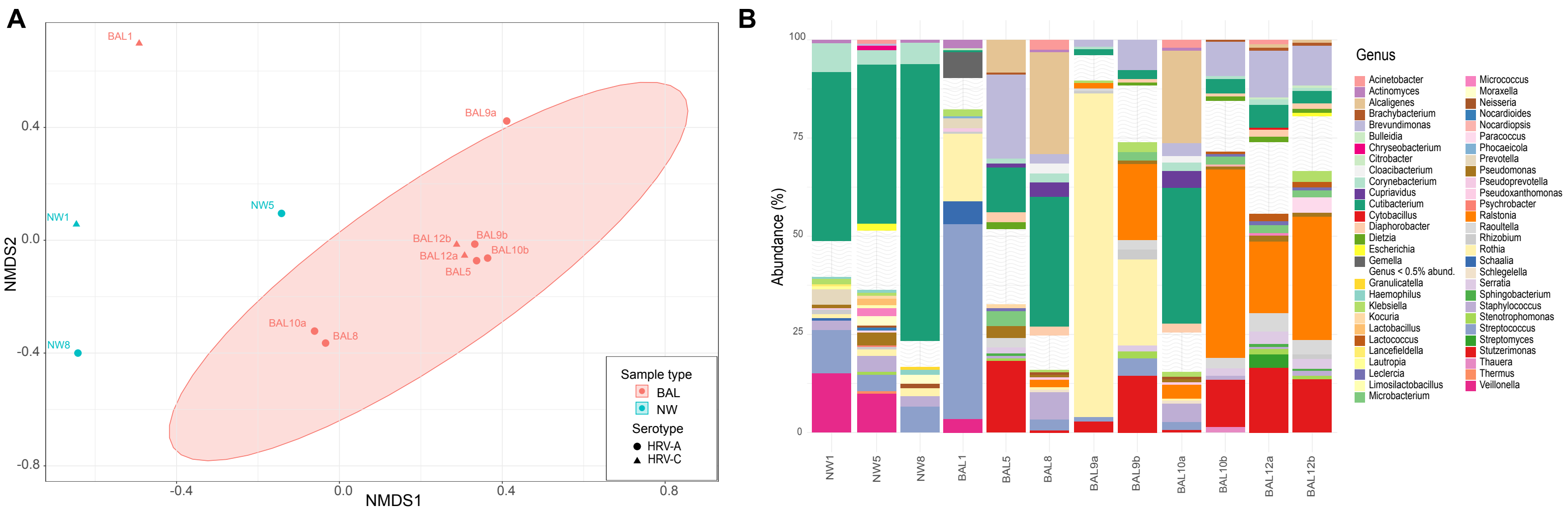

### Supplemental figure 5

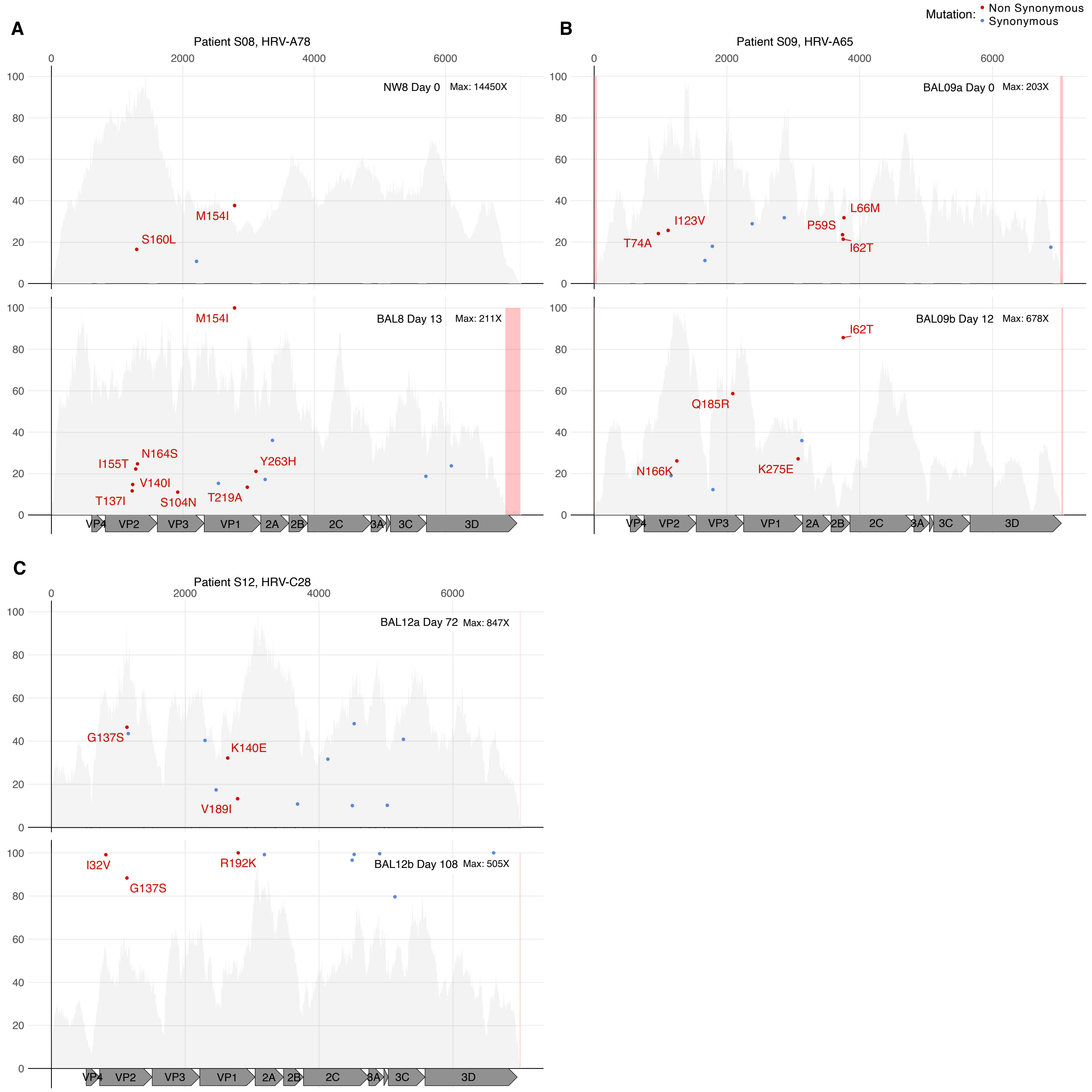

### Supplemental figure 6

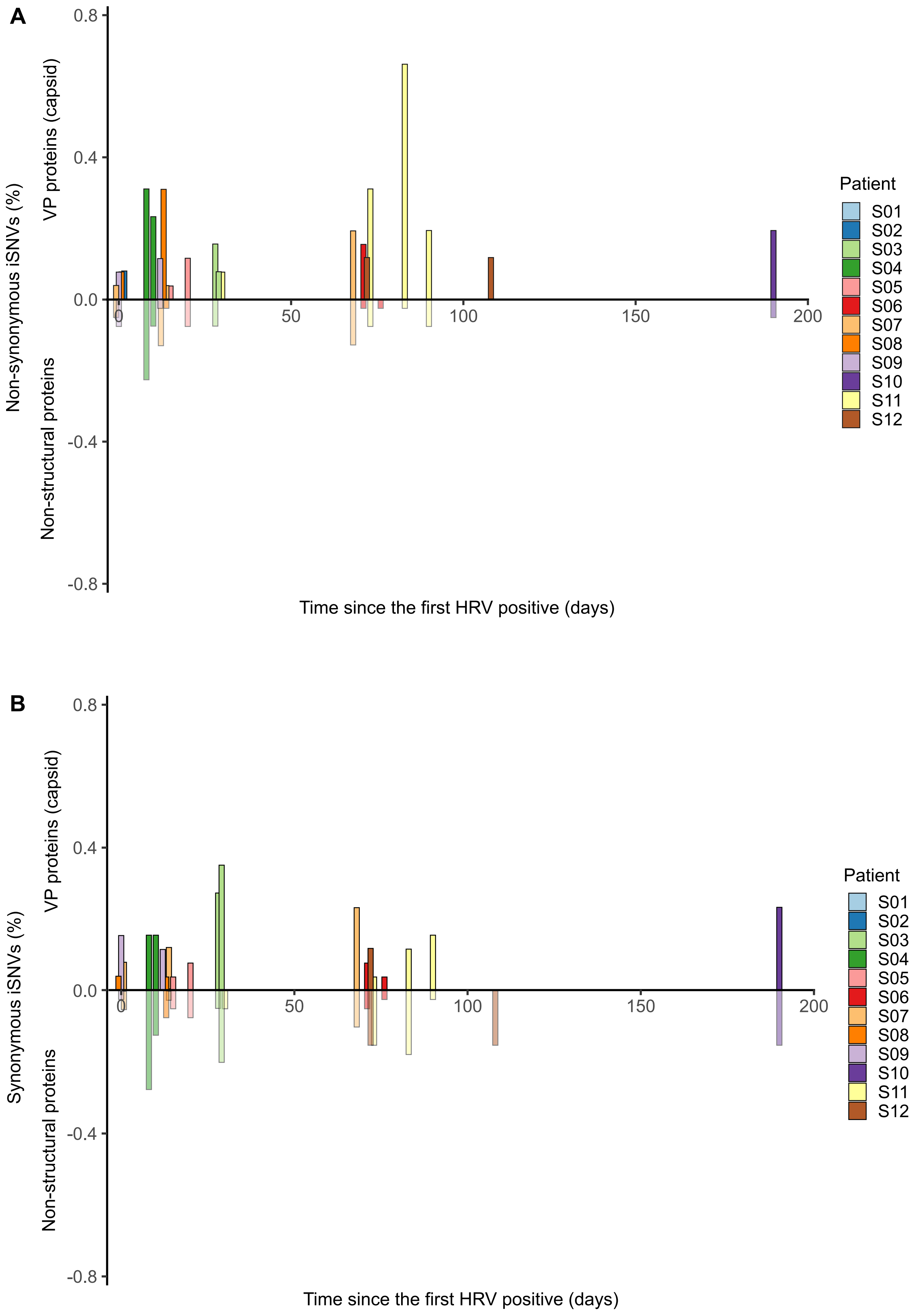

### Supplemental figure 7

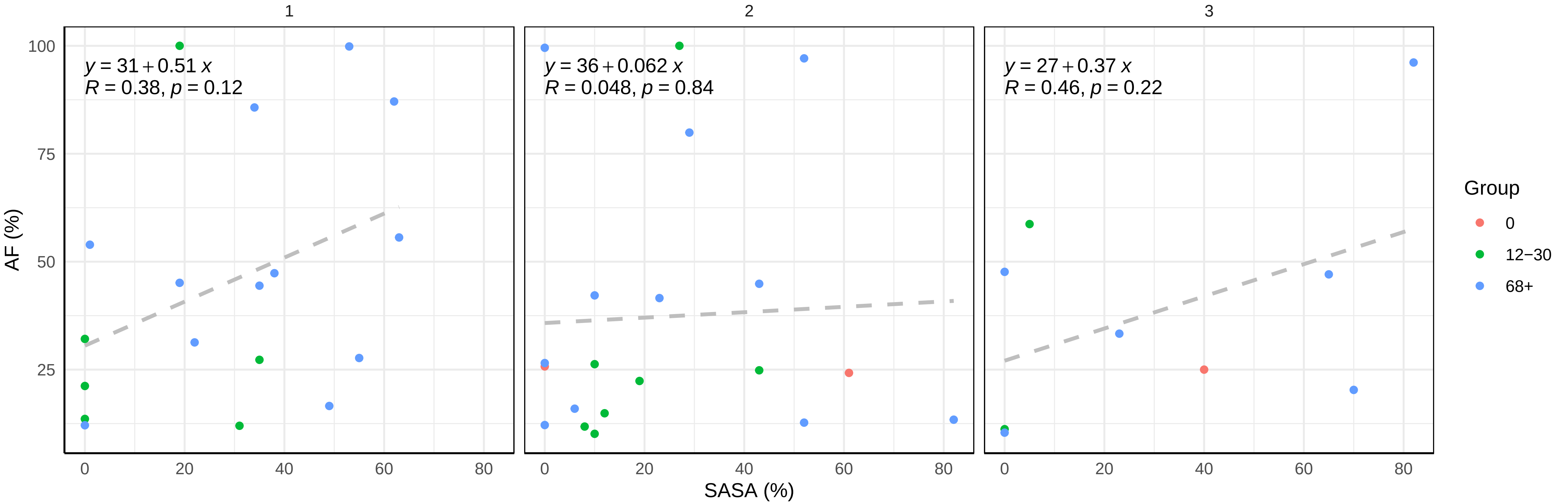

### Supplemental figure 8

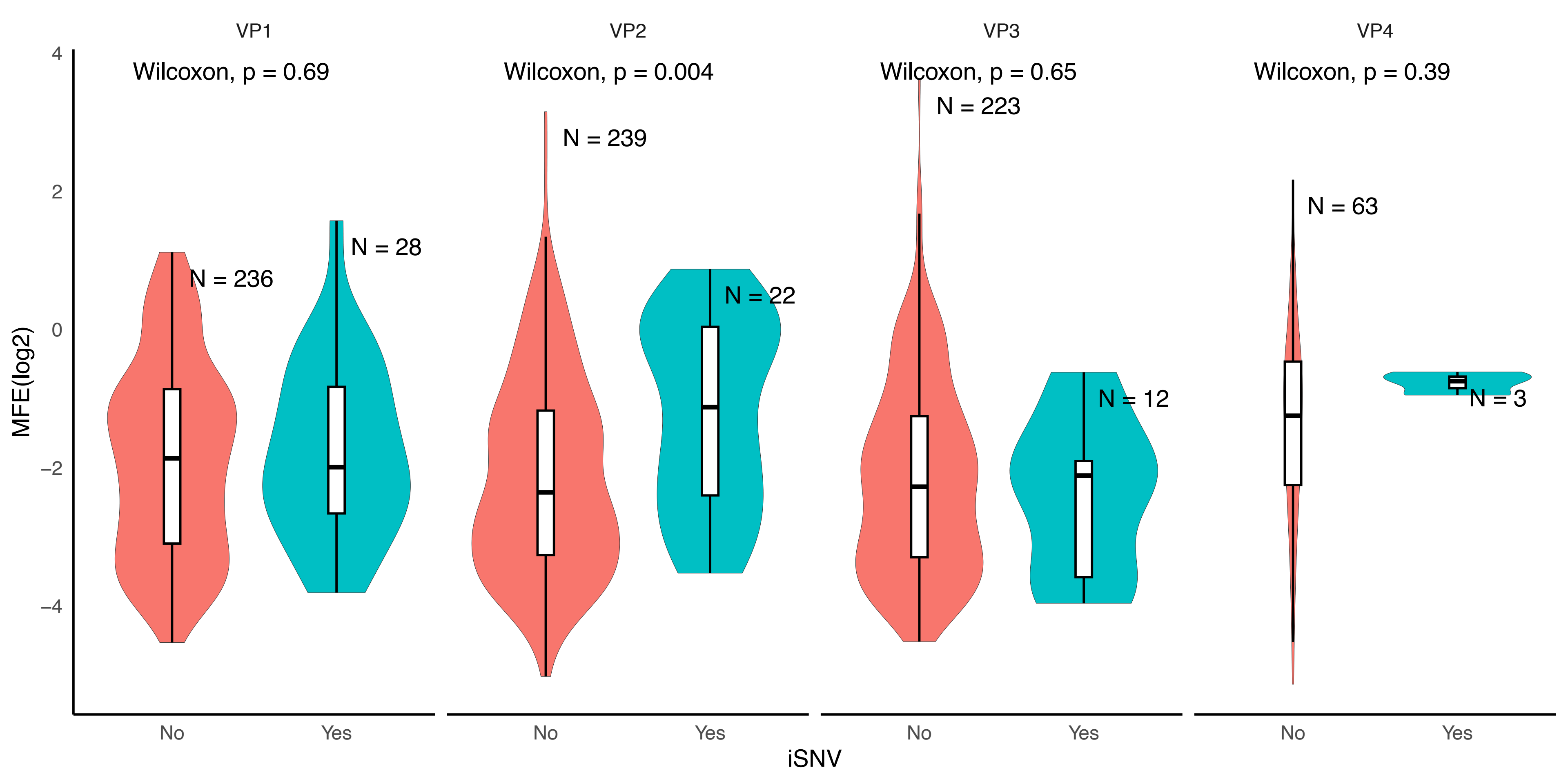

### Supplemental figure 9

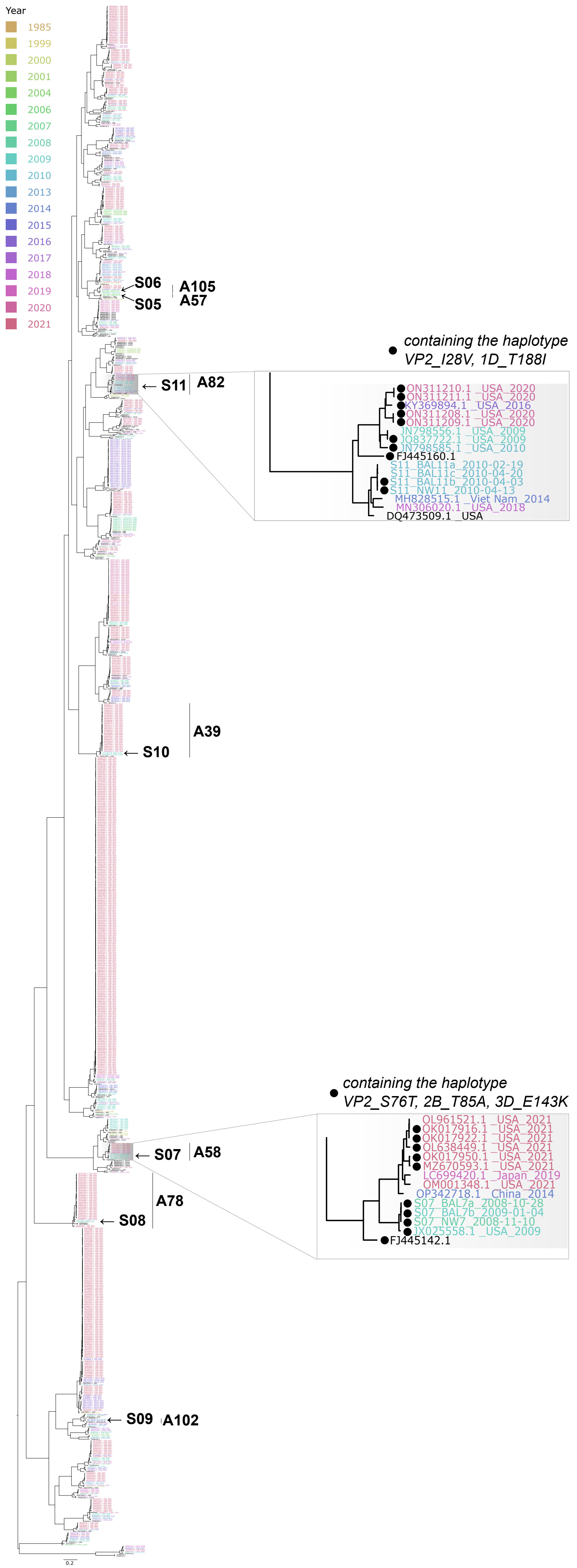
